# Validation of Dual Energy X-ray Absorptiometry for longitudinal quantification of tumor burden in a murine model of pancreatic ductal adenocarcinoma

**DOI:** 10.1101/2023.09.17.558153

**Authors:** Zachary R. Sechrist, Grace Lee, Edward M. Schwarz, Calvin L. Cole

## Abstract

Noninvasive imaging is central to preclinical, in vivo models of pancreatic ductal adenocarcinoma (PDAC). While bioluminescent imaging (BLI) is a gold standard, its signal is dependent on the metabolic activity of tumor cells. In contrast, dual energy X-ray absorptiometry (DEXA) is a direct measure of body composition. Thus, we aimed to assess its potential for longitudinal quantification of tumor burden versus BLI. We utilized the KCKO murine model of PDAC and subjected tumor-bearing (n = 20) and non-tumor control (NTC) (n = 10) animals to weekly BLI and DEXA measurements for up to 10 weeks. While BLI detected tumors at 1-week, it failed to detect tumor growth, displayed a decreasing trend overtime (slope = -9.0x10^8^; p = 0.0028), and terminal signal did not correlate with ex vivo tumor mass (r = 0.01853; p = 0.6286). In contrast, DEXA did not detect elevated changes in abdominal cavity lean mass until week 2 post inoculation and tumors were not visible until week 3, but successfully quantified a tumor growth trend (slope = 0.7322; p<0.0001), and strongly correlated with final tumor mass (r = 0.9351; p<0.0001). These findings support the use of BLI for initial tumor engraftment and persistence but demonstrate the superiority of DEXA for longitudinal tumor burden studies. As tumor detection by DEXA is not restricted to luciferase expressing models, future studies to assess its value in various cancer models and as an in vivo outcome measure of treatment efficacy are warranted.

## Introduction

Pancreatic ductal adenocarcinoma (PDAC) is a highly malignant cancer currently responsible for 8% of cancer related deaths in the United States(1). Despite continued advances and an in-depth understanding of its pathology, the 5-year survival rate remains at a dismal 12%, the lowest among all cancers(1, 2). Preclinical models of PDAC are essential tools that researchers use to investigate mechanisms of tumor initiation, engraftment, and progression, as well as efficacy of novel therapies(3). These studies require reliable in vivo imaging techniques that accurately access tumor growth and response to treatment. While numerous imaging modalities have been utilized for preclinical cancer research, each has its unique strengths and limitations(4, 5).

Bioluminescent imaging (BLI) has been widely adopted as a gold standard due to its relatively low cost and impressive sensitivity(6-8). BLI relies on de novo transgenic expression of firefly luciferase and injection of its luciferin substrate, which produces light that can be detected and quantified as photon/second/cm^2^/steradian (p/s/cm^2^/sr) via a CCD camera system. As such, a major disadvantage of this approach is the requirement of genetically modified tumor models. Furthermore, as BLI signal intensity is a function of tumor metabolism rather than tumor size, it must be used in conjunction with another outcome measure that directly assesses tumor mass(9). Thus far, the focus in preclinical models of PDAC has been ultrasound and magnetic resonance imaging (MRI), which have sensitivity and specificity strengths, but suffer from high costs at low throughput(6). To overcome this, we developed a dual energy X-ray absorptiometry (DEXA) approach to quantify PDAC-induced skeletal muscle wasting using a non-metastatic model of PDAC(10). To expand on this success, we investigated DEXA’s ability to detect the engraftment of orthotopically implanted luciferase expressing PDAC cells (KCKO-Luc), tumor growth over time, and accuracy in predicting ex vivo tumor mass versus BLI, in our murine model.

## Methods and Materials

### Murine orthotopic model of pancreatic cancer

We utilized the murine syngeneic-orthotopic model of PDAC-induced skeletal muscle wasting as described previously by our lab(10), and the overall study design is illustrated in Figure 1. With no previously defined sexual dimorphism in this model, only female C57BL/6J mice were utilized and were obtained from Jackson Laboratory (stock number 000664). Male mice would often fight and reinjure surgical wounds leading to complications therefore, female mice were chosen as the sole model for this research. Mice were maintained in a pathogen-free facility under an Institutional Animal Care and Use Committee’s (IACUC) approved protocol (protocol number: 2018-007). Complete animal protocol is available upon request. All mice were maintained in standard isolation cages with a 12 h light: dark cycle with ad libitum access to water and standard chow. As the development of the KCKO PDAC model has been described elsewhere(11) it will be described in brief as it pertains to this experimentation. Murine KCKO tumor cells were maintained in complete Dulbecco’s modified Eagle’s medium (DMEM) (Corning) supplemented with 10% FBS (HyClone), and 1% penicillin/streptomycin (Gibco). All cell lines were tested negative for mycoplasma. After a 1-week acclimation period, an initial experiment randomized mice to PDAC (n = 10) or non-tumor control (NTC) (n = 10) groups. Prior to inoculation, mice in all groups were subjected to a DEXA scan to establish a baseline measurement. Mice in the PDAC group were anesthetized and injected in the tail of the pancreas with 1x10^5^ KCKO-luc cells suspended in a 1:1 DMEM to Matrigel (Corning) mixture. NTC mice received no surgery. Beginning one-week post tumor inoculation, tumor burden was assessed using both BLI and DEXA imaging. To increase the rigor and reproducibility of this study, an additional 10 PDAC mice were used for manual tumor ROI segmentation studies. As described previously(10, 12), our lab used a murine model of pancreatic cancer to investigate mechanisms related to PDAC-induced SMW. Findings from our lab show that there is no correlation between tumor size and the degree of muscle wasting (data not shown) and that approximately 40% of mice experience a tumor burden larger than the accepted limit of 2000mm^3^ (2g) before the onset of SMW. Our lab recognizes 2 grams as the upper limit of tumor burden based on a conversion from 2000mm^3^ to grams assuming a density of 1g/cm^3^. Therefore, mice that experience a tumor burden larger than 2 grams yet do not experience other characteristics of “failure to thrive” are retained in the study cohort until the required endpoints are met. Additionally, to ensure humane management, specific endpoints were established by us and approved by the University veterinarian and IACUC according to the Guide for the Care and Use of Laboratory Animals (8^th^ edition)(13). In particular, tumor-bearing mice were sacrificed when they developed end-stage disease, defined by exhibiting three or more characteristics of the IACUC definition of “failure to thrive”. Characteristics of “failure to thrive” included but were not limited to, a tumor greater than 2 grams, a loss of lean mass greater than 20% from the baseline DEXA measure, self-isolation, hunched over appearance, lack of or reduced cage activity, lack of or no resistance to scruffing, mangled hair appearance after scruffing, failure to eat or drink, and/or visual signs of breathing difficulty. Mice that displayed tumors greater than the allowed limit only required two additional criteria to be met. To determine if these characteristics were exhibited, animals were checked twice daily, starting 30 days after tumor inoculation as this is the time point in which untreated animals begin to reach advanced-stage disease. When animals were found to exhibit 3 or more characteristics of “failure to thrive” they received a final DEXA scan and were euthanized via carbon dioxide overdose or cardiac puncture and secondary cervical dislocation. Additionally, although rare, KCKO tumor cells may escape the pancreas after injection. This will lead to ectopic growth and in some cases spread towards the thoracic cavity leading to impaired respiration. In these rare events when a mouse experiences atypical tumor growth and hampered breathing it was humanely sacrificed as mentioned without meeting additional criteria. Following euthanasia, tumors were carefully extracted taking care to remove all non-tumor tissue and weighed.

**Fig 1.**
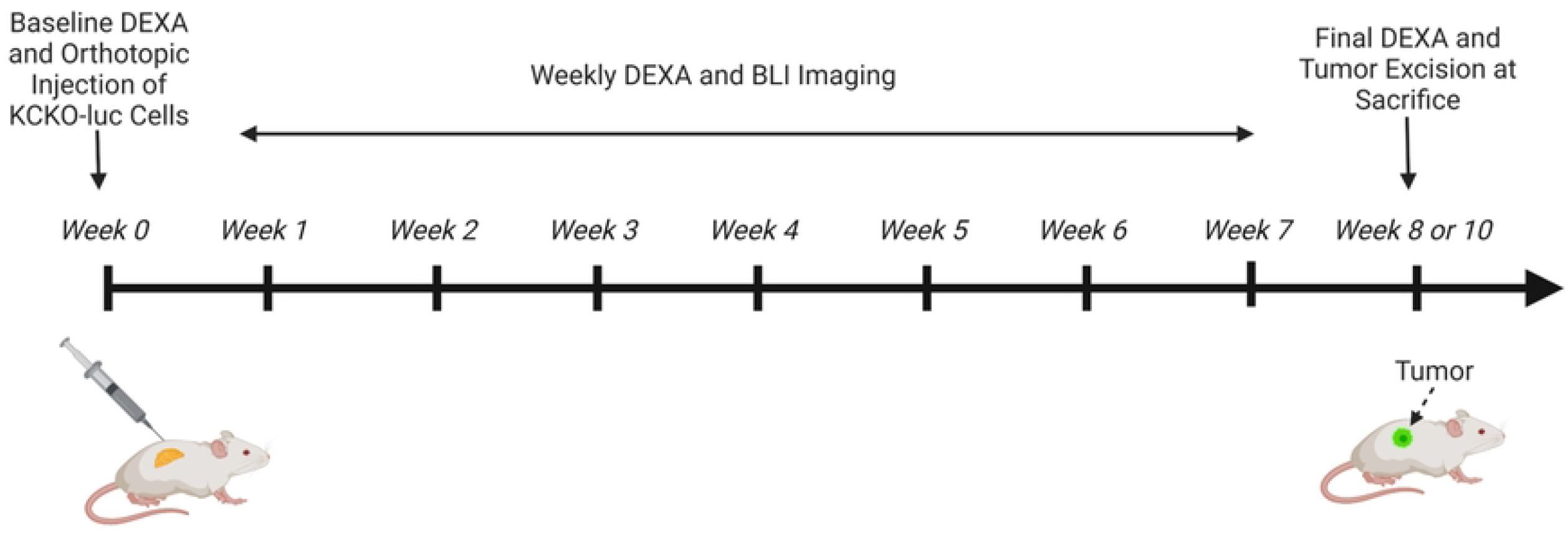
Schematic overview of the orthotopic murine model of PDAC with weekly BLI and DEXA scans. Mice were injected with 1.0x10^5^ KCKO-Luc cells and underwent weekly bioluminescent imaging (BLI) and dual energy X-ray absorptiometry (DEXA) analyses. Mice were monitored for failure to thrive criteria and received a final DEXA measurement before sacrifice on week 8 or week 10 post tumor inoculation. Tumors are excised and weighed postmortem. Created with BioRender.com (2023).

### Bioluminescent imaging

In vivo tumor growth was measured using an IVIS Spectrum Imaging System (IVIS, PerkinElmer). Mice were anaesthetized by vaporized isoflurane and injected subcutaneously (s.c.) with D-luciferin (2.5 mg, Invitrogen) in 100 µl PBS vehicle. While in the right lateral recumbent position, a series of images were taken at 2 min intervals for 24 min, and photon emissions were collected. Bioluminescence (p/s/cm2/sr) was calculated within matching (circular) regions of interest (ROIs) manually placed over tumors. Peak intensity was recorded for each tumor upon two sequential measurements demonstrating signal decay.

### Dual energy X-ray absorptiometry (DEXA)

Body composition was assessed in all mice using a DEXA scanner (iNSiGHT VET DXA; OsteoSys, Seoul, Korea). Mice were weighed before undergoing DEXA scan analysis and anesthetized during imaging via vaporized isoflurane. Each mouse was placed on the scanner bed within the designated scanning area (16.5cm x 25.5cm) in the right lateral recumbent position with the lower limbs stretched away from the abdomen. The Insight DEXA employed a scan time of 30 seconds and utilized a cone beam scanning method generating beams with 60 and 80 kV and 0.8 mA that provided up to a 100-micron resolution for each image. The system provides quantitative data on the bone tissue, fat tissue content, lean tissue content, and the total tissue mass within the region of interest (ROI). Exclusion ROI were used to highlight the skull, ear tag, and nose cone to remove interference as recommended by the manufacturer. The iNSiGHT VET DXA was calibrated daily prior to testing using a quality control phantom according to manufacturer’s instructions. To calculate abdominal lean mass, a scout ROI was drawn defining the region from vertebrae T13-L6 along the spine and extending to the distal end of the femur. Starting 3 weeks post tumor inoculation, a segmented ROI was manually drawn around visible abdominal tumors as identified by lightened pixel color to assess tumor burden more accurately. The lean mass value measured from the segmented ROI from each mouse’s final DEXA scan was used in comparison against ex vivo tumor weights. Additionally, mice were scanned while in the prone position and ROIs were drawn to quantify the lean mass of the lower hindlimbs as described previously(10, 12) (data not shown). Longitudinally monitoring changes in lower limb lean mass ensured that the health of individual mice was accounted for and aided in the determination of humane endpoints.

### Statistical analysis

2-way mixed-model ANOVA tests were used to compare the percent change in abdominal lean mass between NTC and PDAC mice in the weeks post tumor inoculation compared to baseline scans. Simple linear regression models were used to evaluate tumor burden as a function of time post tumor inoculation using BLI and DEXA. Pearson correlation coefficient was calculated to measure the association between BLI measurements of tumor burden and DEXA predicted lean mass versus the weight of the excised tumor upon sacrifice. An intraclass correlation coefficient (ICC) analysis was performed to determine the reproducibility of the DEXA measures to accurately quantify tumor burden from two independent observers. Analyses were performed using GraphPad Prism software (GraphPad Software, San Diego, CA, USA) version 9.5.1. P<0.05 was considered significant (*P<0.05, **P<0.01, ***P<0.001, ****P<0.0001).

### Data Availability Statement

All relevant data are reported within the paper and raw data files are available upon request.

## Results

### BLI is a very sensitive biomarker of KCKO-Luc engraftment but not PDAC mass or growth

To access the reliability and validity of BLI to detect the engraftment and growth of PDAC tumor cells, we used the Muc1-null PDAC model (designated KCKO-Luc) (Figure 1). Longitudinal assessment via BLI following orthotopic implantation of tumor cells into the pancreas of mice confirmed the robust sensitivity to tumor cell engraftment, as strong signal was detected 1 week after implantation (Figure 2). However, no remarkable increase in BLI signal was detected thereafter, and some time points did not have detectable BLI signal, which is consistent with the established paradigm that BLI is primarily a biomarker of KCKO-Luc cell metabolic activity rather than PDAC tumor mass in this model(7, 9).

**Fig 2.**
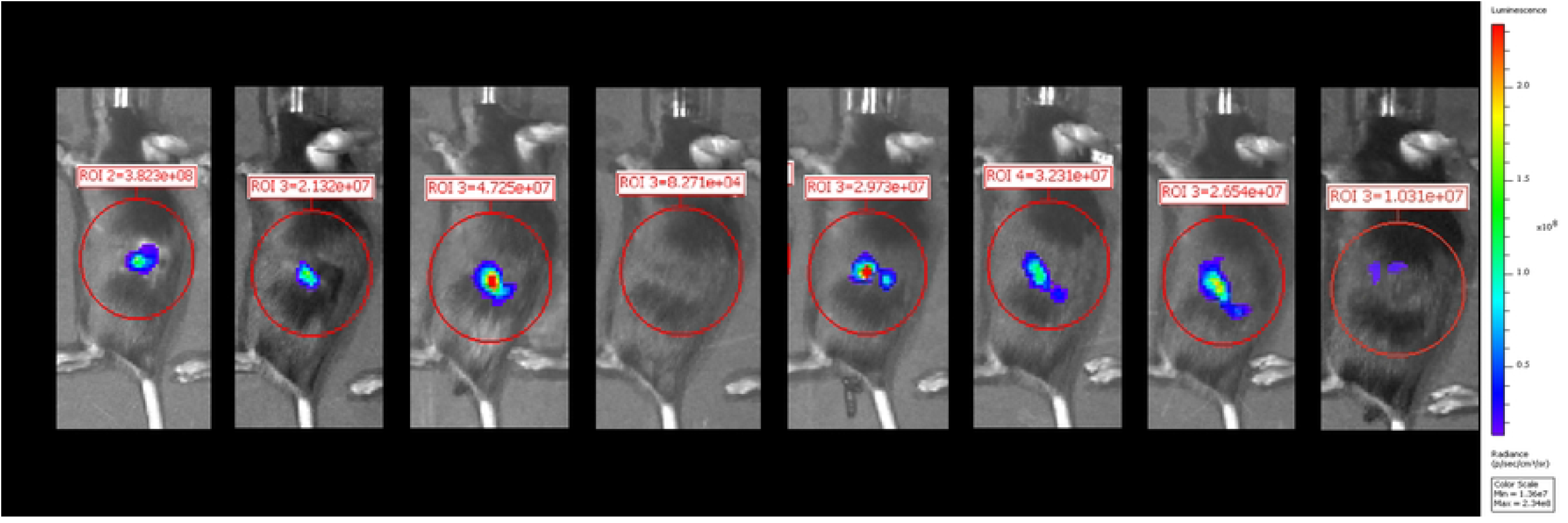
Limitations of bioluminescent imaging as a longitudinal biomarker of tumor mass and growth over time. Longitudinal images of a representative PDAC mouse from week 1 to week 8 post tumor inoculation using bioluminescent imaging (BLI). A standardized abdominal ROI (red circle) was used to quantify BLI values as total flux in radiance (photons/second). Note the robust BLI signal at week 1, the unexplained absence of BLI signal at week 4, and BLI signal at week 8 that is smaller than the BLI signal at week 1.

### Longitudinal quantification of PDAC tumor mass via DEXA

To determine DEXA sensitivity in detecting KCKO tumor burden in vivo, non-tumor control (NTC) (n = 10) and tumor-bearing mice (PDAC) (n = 10) were subjected to weekly DEXA scans from baseline to week 8 post tumor inoculation. Assessment of the scout views revealed dense tumor tissue identified by regions of increased pixel intensity starting at week 3 (Figs. 3A and B). The percent change in DEXA quantified abdominal lean mass for weeks 1 through 8 compared to the baseline shows significant increases in the PDAC mice compared to the NTC animals confirming the presence of a tumor burden starting at week 2 post tumor inoculation (Fig. 3C). An increase in lean mass is detected prior to the visualization of a tumor burden at week 3. Quantification of abdominal lean mass using this approach provided lean mass values that were commensurate with the highlighted tumor burden and ex vivo tumor weights (Figs. 3D and E). These findings confirm the validity of DEXA to identify and longitudinally quantify a growing tumor mass at late timepoints in vivo.

**Fig 3.**
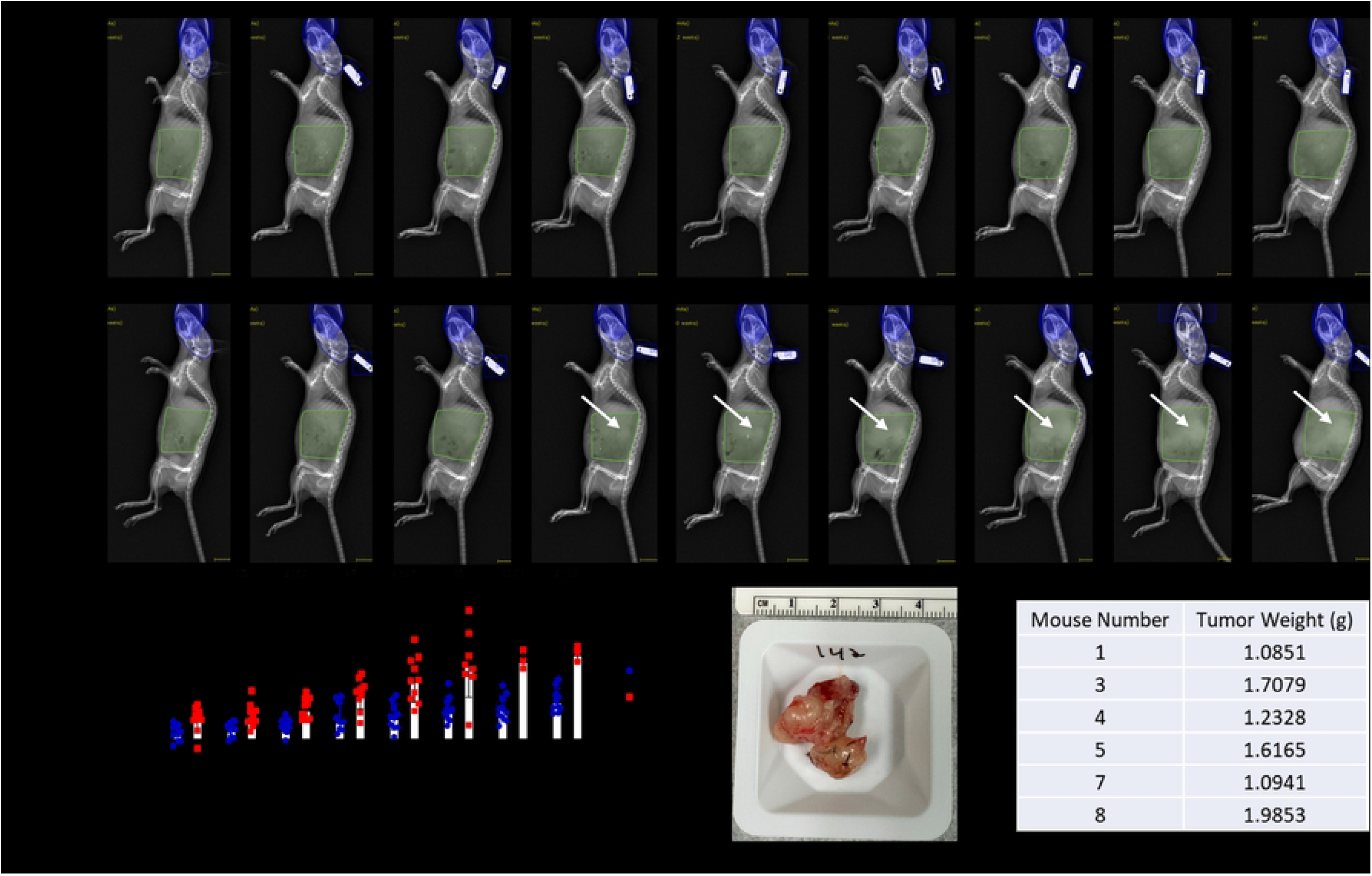
Longitudinal quantification of growing PDAC tumor mass via DEXA and validation with ex vivo tumor weight. Longitudinal DEXA images of a representative non-tumor control (NTC) mouse (A) and PDAC mouse (B) from baseline to week 8 post tumor inoculation are shown. An initial abdominal scout view ROI was used to identify PDAC tumors via manual segmentation from vertebrae T13-L6 and extension to the distal end of the femur (green highlighted region). An exclusion ROI (blue region) was utilized to remove interference from the skull and ear tag. Arrows indicate the presence of the tumor burden visible via DEXA starting at 3-weeks. The weekly percent change in scout ROI abdominal lean mass compared to baseline values from week 1-8 display the significant presence of a tumor burden at week 2 and on (p<0.05) (C). Sample size values represent the living PDAC mice at each weekly DEXA scan. Six PDAC mice met failure to thrive criteria, and their tumors were harvested for gross weight analysis. A representative image of the excised tumor from the median PDAC mouse weighing 1.6165 grams and 3-4 cm^2^ is shown (D), with all the ex vivo tumor weights from the PDAC mice that were not lost to attrition (E).

### Validation of DEXA as a reliable tool to longitudinally assess tumor growth

To further establish the advantages of DEXA over BLI for measuring tumor burden in vivo, ten additional PDAC mice were scanned weekly from baseline to week 10 post tumor inoculation and longitudinal outcomes of BLI signal and abdominal lean mass were compared (Figure 4). Attempts to quantify PDAC burden from previously mentioned scout ROI failed due to the great variability in tumor morphology (data not shown) and interference from non-tumor tissue. Thus, we adopted a manual tumor focused ROI segmentation (Fig. 4A) which was indicative of the additional ex vivo tumor weights (data not shown). The results showed that DEXA measurements were consistent in individual mice over time, and the average lean mass measures for all PDAC mice in the study (n = 20) demonstrated a significant increasing trend over time (slope = 0.7322, p<0.0001) (Figs. 4B and C). In contrast, BLI was not able to detect the increasing tumor burden over time, perhaps due to a ceiling effect as the maximal signals in our study were detected 1-week post implantation of the KCKO-Luc cells. Discordance between BLI signal and tumor mass was further illustrated by continuous weekly fluctuation in individual mice with PDAC (n = 10), as well as undetectable signal (total flux<10^6^ p/s) at some time points. Moreover, the average BLI signal for all tumor-bearing mice (n = 20) decreased over time (slope = -9.0x10^8^, p = 0.0076) (Figs. 4D and E), which is inconsistent with the observed tumor growth. Additionally, direct assessment of the biomarkers versus ex vivo tumor weight confirmed no correlation between the terminal BLI signal and PDAC mass (Fig. 4F). In contrast, DEXA measured lean mass proved to be a very strong predictor of tumor weight (Fig. 4G). Lastly, segmented ROI demonstrates excellent interobserver reproducibility (ICC = 0.9736) making it a reliable analysis tool for longitudinal tumor studies (Fig. 4H).

**Fig 4.**
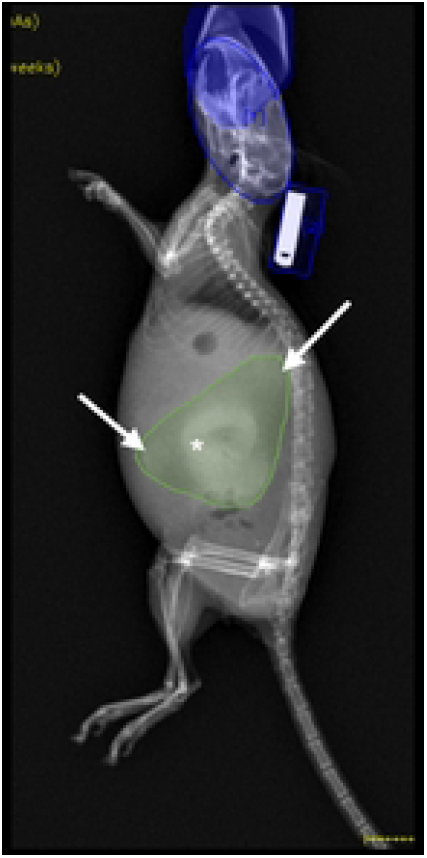
Superiority of DEXA over BLI as an in vivo biomarker of tumor mass and longitudinal measure of tumor growth. Ten additional mice received orthotopic KCKO-Luc cells and the growth of their PDAC was assessed via BLI and DEXA until their failure to thrive or sacrifice at 10 weeks post-implantation. DEXA quantification for this study was derived from manual ROI segmentation which reports the focused PDAC ROI in grams (A). The asterisk denotes the primary tumor and arrows indicate local spread. Longitudinal DEXA quantification from each mouse is presented starting when tumor burden is visible (B) as well as the mean +/-SD and slope for all PDAC mice (n = 20) from weeks 3-10 (C). Weekly BLI signal is presented for each mouse in this study (n = 10) (D) as well as the mean +/-SD for all tumor-bearing mice (n = 20) (E) from weeks 1-10 with the slope. To assess the correlation between PDAC mass and biomarker outcome, linear regression analyses were performed comparing ex vivo tumor weight vs. terminal BLI (F) and terminal DEXA measures (G), and the graphed data are presented with the Pearson coefficient and p-value. Two of the authors completed independent focused PDAC segmentation on end-of-life DEXA scans, and the interobserver reliability of the quantified tumor mass was determined via intraclass correlation coefficient (ICC r^2^ = 0.9736; p<0.0001) (H). n = 20 week 1-5, n = 16 week 6, n = 11 week 7, n = 7 week 8, n = 6 week 9, n = 5 week 10.

## Discussion

To date, there have been numerous imaging modalities tested and validated for the detection and longitudinal assessment of in vivo tumors(4, 5). It is understood that no imaging technique exists without limitations(4, 14, 15). Therefore, it is essential that the researcher fully understands their model and utilizes a combination of methods. This is particularly true when investigating debilitating diseases such as PDAC. As our lab is focused on skeletal muscle wasting as a result of late-stage PDAC, our murine model is subjected to weekly BLI and DEXA measurements as described by our lab previously(10, 12). We have observed that increased mouse handling and constant anesthetization further exacerbates a decline in health. Thus, the present study aims to adapt DEXA for the reliable and reproducible quantification of tumor burden in an effort to reduce the stress placed on these animals.

We demonstrated the reliability of DEXA to detect tumors starting at week 3 post inoculation. Beginning at week 2, tumor growth was detectable compared to healthy controls. A decided advantage of DEXA is the quantitative readout of tumor size in grams which displays a strong correlation to the weights of tumors resected from mice. Conversely, BLI reports tumor burden in arbitrary units making it an unreliable measure of tumor volume. Furthermore, because BLI requires de novo luciferase protein synthesis and ATP breakdown, only metabolically active cells produce detectable levels of bioluminescence(7). This was demonstrated in our study as tumor induced luminescence reaches its maximum intensity one week after inoculation, and slowly declines as tumors become dense and necrotic. Fortunately, as the tumor continuously develops a denser core, DEXA is able to identify and longitudinally monitor growth. It should be noted that individual tumors are detectable on DEXA at different time points with the majority being visible at 3-weeks. This can be explained by the heterogeneous nature of PDAC and variability in tumor density, ultimately determined by the vascular network(16) and varying rates of engraftment. Furthermore, defining the tumor ROI on DEXA is reliant on the presence of lightly colored dense tissue and is therefore, open to subjectivity in analysis. Variation in tumor quantification may arise for tumors with local secondary masses of lowered density. Fortunately, the growth rate of the KCKO tumor model is comparatively slow for PDAC and frequent weekly DEXA scans accurately identify the growing tissue. Moreover, at late timepoints, the tumor often invades surrounding structures such as the spine, kidneys, and spleen. Incomplete removal may be an additional discrepancy in ex vivo tumor weight. Despite these minor limitations, we show that DEXA quantification correlates strongly with ex vivo tumor weight and provides reproducible measures when analyzed by independent observers, validating it as a reliable method for longitudinal tumor quantification.

A major limitation of BLI analysis is the dependency on luciferase reporters. PDAC research is widely reliant on genetically engineered and patient derived xenograft (PDX) models that better recapitulate aspects of human disease(17, 18). As such, they cannot readily incorporate luciferase reporters as done with injectable models. It then becomes necessary to utilize other imaging techniques. Ultrasound and MRI are commonly used for tumor detection but require immense time and resources, and require add-on software like Amira for tumor quantification(19, 20). Based on our study in an orthotopic model, DEXA has the potential to be used in conjunction with ultrasound and MRI as a cost effective and time efficient determinant of tumor size in these additional PDAC models and requires further investigation.

The sensitivity in which BLI detects tumor engraftment at early timepoints makes it ideal for validation of tumor engraftment. In our prior research with xenograft models of cancer in bone, in which tumor engraftment efficiency is ∼50% at 3 weeks, we found BLI to be indispensable for exclusion of non-tumor bearing mice and randomization to treatments (21, 22). However, in these studies we also noted that BLI signal does not correlate with tumor size, vascularity and osteolysis. Consistently, we also found here that BLI signal decreases over time and is a poor indicator of tumor size. Based on this, it is proposed that BLI be used at early time points in our model with a reliance on DEXA for accurate quantification of tumor size at late timepoints. Additionally, DEXA enhances the use of tumor burden as a failure to thrive criteria in our model. Adhering to an experimentally derived threshold of tumor size will inform when mice reach a humane endpoint and will ultimately improve living conditions. Furthermore, the use of DEXA as a dual method to assess SMW and tumor burden will reduce the need for additional bouts of sedation which further comprises animal health during the late stages of disease. Lastly, it is speculated that DEXA will be a reliable tool to identify the efficacy of anti-tumor therapies in vivo across a variety of models. DEXA will continue to be a multipurpose tool in our lab and aid in the understanding of tumor growth and PDAC induced SMW.

## Acknowledgements

This work was supported by grants from the National Institutes of Health, K01 CA240533. We thank the members of the Histology, Biochemistry & Molecular Imaging Core and the Biomechanics, Biomaterials, and Multimodal Tissue Imaging Core in the Center for Musculoskeletal Research.

